# Loss of PREPL alters lipid homeostasis due to mitochondrial defects

**DOI:** 10.1101/2025.10.28.685080

**Authors:** Yenthe Monnens, Kritika Bhalla, Karen Rosier, Rita Derua, Anne Rochtus, Celien Lismont, Johan Swinnen, Marc Fransen, John W.M. Creemers

**Affiliations:** Laboratory for Biochemical Neuroendocrinology, Department of Human Genetics, KU Leuven, Leuven, Belgium; SyBioMa, Proteomics Core Facility KU Leuven, Leuven, Belgium; Department of Pediatrics, Section Pediatric Endocrinology, University Hospital Leuven, Leuven, Belgium; Laboratory of Peroxisome Biology and Intracellular Communication, Department of Cellular and Molecular Medicine, KU Leuven, Leuven, Belgium; Laboratory of Lipid Metabolism and Cancer, Department of Oncology, KU Leuven, Leuven, Belgium

## Abstract

Loss of the prolyl endopeptidase-like (PREPL) protein causes congenital myasthenic syndrome-22 (CMS22), a rare neuromuscular and metabolic disorder. PREPL belongs to the serine hydrolase superfamily, but its physiological substrates remain unknown. Based on the predicted lipid binding pocket in its crystal structure and its *in vitro* esterase activity, we hypothesized that PREPL might act as a lipase *in vivo* and directly regulate lipid metabolism. To test this, we performed unbiased lipidomics in *Prepl* knockout (KO) mouse brains and CRISPR-Cas9-generated KO cell lines. Across tissue and cell types, global phospholipid composition was largely unchanged, with only modest, non-significant increases in lysophospholipids, arguing against a direct role of PREPL in (lyso)phospholipid turnover. In contrast, *PREPL* KO HEK293T cells exhibited a significant accumulation of triacylglycerols (TAGs) and an increased number of lipid droplets, indicating a selective shift toward lipid storage. Given the central role of peroxisomes in lipid metabolism, we assessed PREPL localization and examined peroxisome number, morphology, and levels of key peroxisomal proteins. PREPL did not localize to peroxisomes, and peroxisome number and proteins levels were largely unchanged. However, KO cells displayed elongated peroxisomes, a phenotype possibly linked to mitochondrial dysfunction. Indeed, previous studies have shown that PREPL localizes to mitochondria and is required for respiratory chain activity and oxidative phosphorylation. These mitochondrial defects are predicted to impair fatty acid β-oxidation and disrupt redox balance, thereby promoting TAG synthesis and lipid droplet biogenesis as adaptive responses. Overall, our findings indicate that PREPL does not act as a canonical lipase but indirectly alters lipid homeostasis through its critical role in mitochondrial function. Elevated TAG levels and altered peroxisome morphology likely represent secondary consequences of impaired mitochondrial fatty acid metabolism in PREPL-deficient cells. These results establish a mechanistic link between mitochondrial dysfunction and lipid remodeling in PREPL deficiency, providing novel insights into the metabolic pathology of CMS22.

## Introduction

Mutations in *PREPL*, a gene on chromosome 2, lead to the rare autosomal recessive disorder congenital myasthenic syndrome-22 (CMS22) (1–6). Clinically, CMS22 manifests with severe neonatal hypotonia, poor suck, failure to thrive, and growth retardation. As patients grow older, additional features often emerge, including impaired cognitive development, hyperphagia, and obesity. Together, these symptoms indicate that PREPL plays a critical role in both neuromuscular and metabolic physiology. PREPL belongs to the serine hydrolase superfamily and has been implicated in the regulation of the secretory pathway and mitochondrial respiration (4,5,7–9). PREPL has its highest expression in the brain, followed by intermediate levels in neuroendocrine cells, kidney and muscle. Recent studies have reported altered mitochondrial morphology, reduced cristae density, impaired respiratory chain activity, and compromised oxidative phosphorylation in *PREPL* knockout (KO) cells and tissues. While part of the functions of PREPL appear to be regulated through non-catalytically protein-protein interactions, catalytic activity remains crucial for mitochondrial respiration. Despite these insights, the precise molecular mechanisms by which PREPL deficiency leads to disease remain elusive. In particular, the identity of PREPL’s physiological substrates has yet to be established, making substrate discovery essential for understanding its function and the pathophysiology of CMS22.

Unlike its close homologue prolyl endopeptidase (PREP), PREPL does not exhibit peptidase activity (10,11). Instead, recent structural and biochemical findings point toward alternative enzymatic roles. The crystal structure of PREPL revealed an α/β-hydrolase fold architecture coupled with a β-propeller domain, a combination often associated with substrate selectivity in hydrolases. Activity-based probe profiling showed that PREPL can be competitively inhibited by Palmostatin M, a well-characterized inhibitor of the acyl protein thioesterases APT1 and APT2 (4). In line with this observation, PREPL was found to hydrolyze ester and thioester substrates *in vitro* (4). Docking experiments further identified a putative lipid-binding pocket within the catalytic domain, suggesting that lipids could represent natural substrates. Lipids such as triacylglycerols are composed of glycerol and fatty acid chains connected by ester bonds. Therefore, we investigated if PREPL possesses lipase activity *in vivo* and whether impaired lipid metabolism is the central pathogenic mechanism underlying CMS22.

## Materials and methods

### Lipidomics of mouse brain and β-TC3 cells

*Prepl* mice were sacrificed via cervical dislocation and transcardially perfused with ice-cold PBS. Following dissection of the brain, the hemispheres were separated, quickly frozen in liquid nitrogen, and stored at −80°C until further use. Tissue preparation and analysis of (lyso)phospholipid species was performed by electrospray ionization tandem mass spectrometry on a hybrid triple quadrupole/linear ion trap mass spectrometer as previously described (12). Prior to quantification of specific lipid species, raw data were subjected to background subtraction, isotope correction, and fatty acid chain length correction (for phospholipids). As background, the intensities of species detected in ‘internal standards only’ spectra were divided by the ion suppression factor of each sample. The ion suppression factor was calculated for each sample separately by dividing the intensity of the standards in the ‘internal standards only’ spectrum by the intensity of the standards in the sample spectrum. Data was normalized based on the amount of DNA. To quantify the total amount of lipids in a specific lipid class, the abundances of individually measured species within this lipid class were summed.

### Lipidomics of HEK293T cells

Lipidomics of HEK293T cells was performed as described (13). In brief, lipid extraction was performed on *PREPL*-/- and control HEK293T cells. Sample was homogenized in 700 μL of water and then mixed with 800 μl of 1 M HCl and CH_3_OH in a ratio of 1:8 (v/v), followed by the addition of 900 μl CHCl_3_, 200 μg/ml antioxidant 2,6-di-tert-butyl-4-methylphenol (Sigma Aldrich), and 3 μl SPLASHR LIPIDOMIXR Mass Spec Standard (Avanti Polar Lipids). After centrifugation, the lower organic layer was collected, and the solvent was evaporated using a Speedvac spd111v (Thermo Fisher Scientific) at room temperature. The resulting lipid pellets were reconstituted in 100% ethanol for mass spectrometry analysis. The pellets were analyzed using liquid chromatography-electrospray ionization tandem mass spectrometry (LC-ESI/MS/MS) on a Nexera X2 UHPLC system (Shimadzu) coupled with a hybrid triple quadrupole/linear ion trap mass spectrometer (6500+ QTRAP system; AB SCIEX). Chromatographic separation was performed on an XBridge amide column. Lipids were quantified by multiple reactions monitoring (MRM), and data analysis was conducted using MultiQuantTM software version 3.0.3. To ensure accuracy, the signals of lipid species were adjusted to account for isotopic contributions, calculated using Python Molmass 2019.1.1. Quantification was based on internal standard signals and followed the guidelines outlined by the Lipidomics Standards Initiative (LSI). Unpaired t-test, p-values, and FDR-corrected p-values (using the Benjamini/Hochberg procedure) were calculated in Python StatsModels version 0.10.1.

### Lipid droplet staining and high-content microscopy

*PREPL*^-/-^ and control HEK293T cells were plated on poly-D-lysine (Merck Millipore) coated 96-well Cell Carrier Ultra microplate (PerkinElmer) with a cell density of 15,000 cells/well. The cells were washed thrice with DPBS (pH 7.4; Thermo Fisher Scientific) and fixed using 4% paraformaldehyde (PFA) for 30 minutes. Lipid droplet staining was performed according to the manufacturer’s protocol. In brief, LipidSpot 488 (Biotium) stain was diluted to 1X in DPBS, and cells were stained in solution for 30 minutes in a light-protected environment. This was followed by DAPI (nuclear) staining for 10-20 minutes. Fluorescence was measured using a high-content screening microscope, Operetta (CLS+Twister; PerkinElmer), and lipid droplet size and number were quantified using Harmony 4.9 software (PerkinElmer).

### Peroxisome biology

Immunofluorescence – Staining of individual peroxisomes, as well as co-staining of PREPL and peroxisomes, was performed using the PEX14 (14) and PREPL abnova Maxpab (H00009581-B01P) (15) antibodies, with Alexa-conjugated secondary fluorophores. Images were acquired using a Nikon C2 confocal microscope and analyzed with ImageJ software. Western blotting – Protein concentrations were determined using the Pierce™ BCA Protein Assay kit (Thermo Fisher Scientific). Proteins were denatured, separated by SDS-PAGE on a 10% Bis-Tris gel, and transferred to a nitrocellulose membrane. Membranes were blocked with 0.1 M Tris-HCl (pH 7.4) and 0.15 M NaCl containing 0.5% blocking reagent (Roche) and 0.2% Triton X-100. Membranes were incubated with primary antibodies for 1 hour at room temperature or overnight at 4°C, followed by HRP-conjugated secondary antibodies (1:3000) (Dako). Detection was performed using the Western Lightning ECL system (Perkin Elmer). The antibodies used were directed against GAPDH (Cell Signaling Technology, 2118), PEX14 (15), catalase (Calbiochem, 219010), ACAA1 (Atlas Antibodies, HPA007244), and HSD17B4 (Fisher Scientific, 15116-1-AP). Western blots were quantified using ImageJ software and results were analyzed by T-test using the Graphpad Prism software 8.3.0.

## Results

### Lipidomics of *Prepl* KO brains reveal that PREPL is not involved in (lyso)phospholipid metabolism

To investigate whether PREPL has lipase activity, we first performed lipidomic profiling from *Prepl* KO mice tissue. Since PREPL is highest expressed in the brain, brain samples were obtained and analysed. Untargeted lipidomic profiling of brain tissue revealed the presence of major phospho- and sphingolipid classes, including phosphatidylcholine (PC), phosphatidylethanolamine (PE), phosphatidylinositol (PI), phosphatidylserine (PS), sphingomyelin (SM), ceramide (Cer), as well as their corresponding lysophospholipids (lysoPC, lysoPE, lysoPI, lysoPS). No significant differences were detected in these lipid compositions between wild-type (WT) and KO brains. A non-significant trend toward increased lysophospholipids was observed, with KO brains showing approximately 12% higher abundance across several classes (lysoPC +9%, lysoPE +17%, lysoPI +13%, lysoPS +11%) (Fig 1A-B). These findings raised the possibility that PREPL might participate in lysophospholipid metabolism, similar to what has been described for APT1 (16).

**Figure 1.**
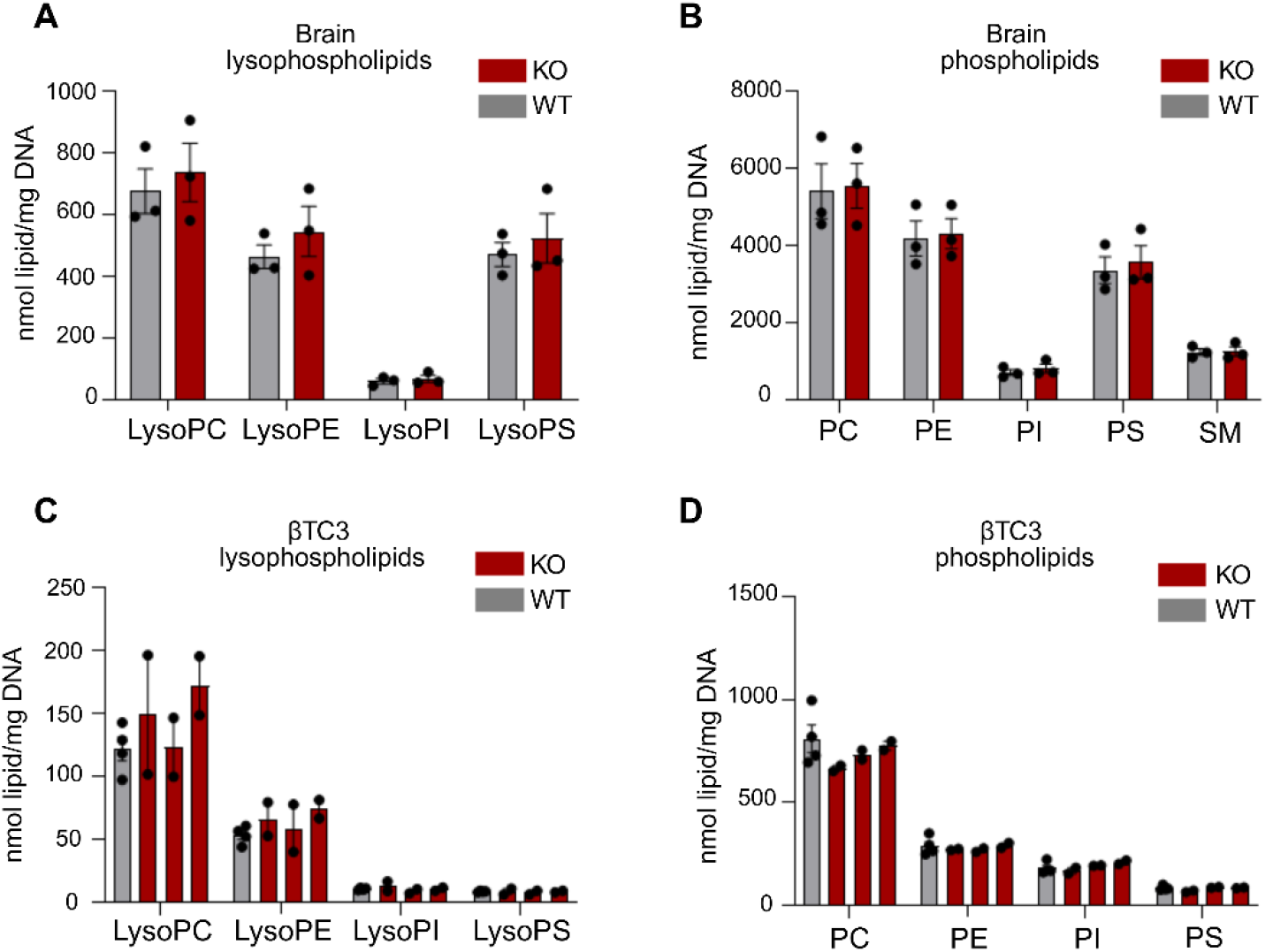
Lipid profile of brain and βTC3 cells are unaltered upon *Prepl* KO. A-B semiquantitative analysis of lysophospholipids (A) and phospholipids (B) in hemisphere of 9-month old mice (n=3) C-D. Lysophospholipid (C) and phospholipid (D) profile from WT and 3 different CRISPR-Cas9 generated *Prepl* KO βTC3 cell lines (n=2-4). Phosphatidycholine (PC), phosphatidylinositol (PI), phosphatidylethanolamine (PE), phosphatidylserine (PS, sphingomyelin (SM).

Since brain is composed of many different cell types, changes in a single cell type may be difficult to detect. Therefore, we analyzed the lipid profile of CRISPR-Cas9-generated *Prepl* KO βTC3 mouse insulinoma cells and detected major lipid classes, including phosphatidylcholine (PC), phosphatidylethanolamine (PE), phosphatidylinositol (PI), phosphatidylserine (PS), sphingomyelin (SM), ceramide (Cer), and their lysophospholipid counterparts (lysoPC, lysoPE, lysoPI, lysoPS). Again, we observed a trend toward elevated lysophospholipid levels, in particular lysoPE (+21%) and lysoPC (+24%), compared with WT controls (Fig 1C-D). However, these differences did not reach statistical significance, and high variability was observed among biological replicates. At the level of individual lipid species, no consistent alterations were found across all samples.

Taken together, these results indicate that while PREPL deficiency may subtly influence lysophospholipid abundance, the evidence does not support a direct role for PREPL in (lyso)phospholipid metabolism.

### *PREPL* KO leads to an increase in triacylglycerols in HEK293T cells

Untargeted lipidomic profiling of WT and *PREPL* KO HEK293T cells detected a broad spectrum of lipid classes, including sphingomyelins (SM), ceramides (Cer), hexosylceramides (HexCer), cholesteryl esters (CE), diacylglycerols (DG), triacylglycerols (TG), phosphatidylcholines (PC, PC-O, PC-P), lysophosphatidylcholines (LPC), phosphatidylethanolamines (PE, PE-O, PE-P), lysophosphatidylethanolamines (LPE), phosphatidylglycerols (PG), phosphatidylinositols (PI), and phosphatidylserines (PS), each represented by multiple fatty acyl species ranging from C14 to C26 with varying degrees of saturation (Fig 2). Similar to the results above, most lipids remain unaffected. However, a significant increase in triacylglycerols (TAGs) was observed in *PREPL* KO HEK293T cells. Specifically, triacylglycerol 16:0, 18:1, and 18:3 are significantly increased (Fig 2A). Phosphatidylcholines (PC), phosphatidylethanolamines (PE) and sphingomyelins (SM) remained unaffected (Fig 2B-D). Since triacylglycerols are the main lipids stored in lipid droplets, we evaluated lipid droplet size and quantity in *PREPL* KO HEK293T cells (Fig 2E-F). Consistent with the lipid profile, we found an increase in the number of lipid droplets in KO (14.85 ± 1.11) compared to WT (8.84 ± 0.58) cells.

**Figure 2.**
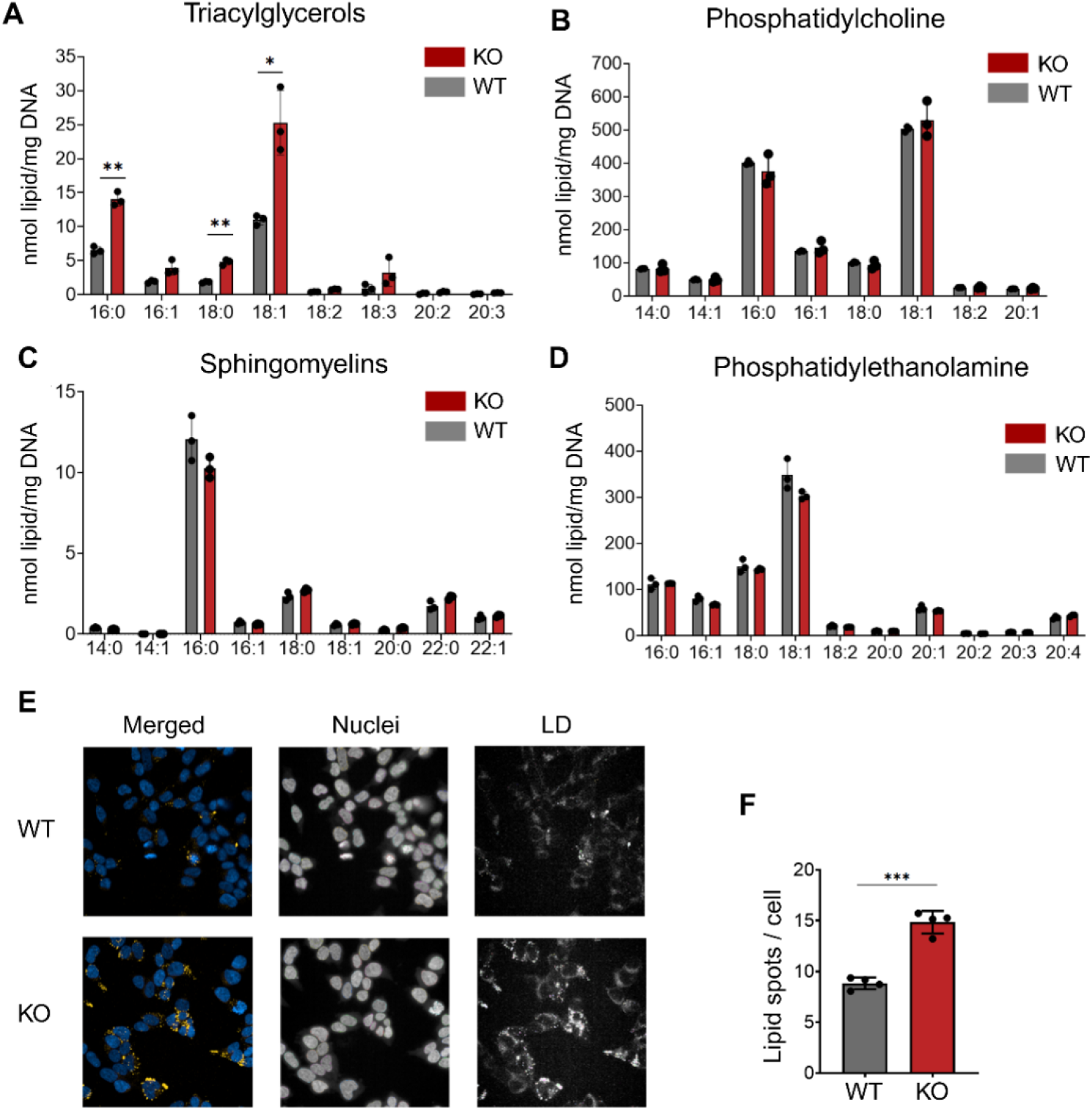
*PREPL* KO leads to an increase in triacylglycerol storage in HEK293T cells. (A) increased levels of fatty acid chains of TAGs in *PREPL* knockout HEK293T cells compared to WT. (B-D) unaltered fatty acid chain content in phospholipids including phosphatidylcholine, sphingomyelins, and phosphatidylethanolamine. (E) Lipid droplet staining of *PREPL* KO and WT HEK293T cells. (F) Quantification of lipid spot numbers/cell, total of 4 independent experiments. ^*^p < 0.05, ^**^p < 0.01, ^***^p < 0.001

### PREPL is not involved in peroxisome biology

Peroxisomes cooperate with mitochondria and lipid droplets to regulate fatty acid oxidation, remodeling of complex lipids, and energy homeostasis(17). Peroxisomes are known to harbor esterase and thioesterase enzymes that hydrolyse fatty acids. Therefore, we hypothesized that PREPL may have a lipid-metabolizing role within peroxisomes. By confocal microscopy, we found that PREPL does not colocalize with the peroxisomal marker PEX14 (Fig 3A). Moreover, Very-long-chain fatty acids (VLCFAs) were not significantly altered in our lipidomic data, suggesting preserved peroxisomal function.Consistent with the lipidomic results, we therefore state that PREPL is not involved in peroxisomal lipid metabolism. However, we observed that loss of PREPL alters peroxisome morphology in *PREPL* KO conditions (Fig 3B). Specifically, peroxisomes are significantly longer in *PREPL* KO cells (1341 ± 360) than in WT cells (1068 ± 291), while the total number of peroxisomes remains unaltered (Fig 3C). Nevertheless, the expression levels of well-established peroxisomal markers – PEX14 (membrane marker), HSD17B4 (matrix protein with a typical C-terminal targeting signal), ACAA1 (matrix protein with an N-terminal targeting signal), and catalase (matrix protein with an atypical C-terminal targeting signal) – remain unchanged in *PREPL* KO cells compared to WT cells (Fig 3D).

**Figure 3.**
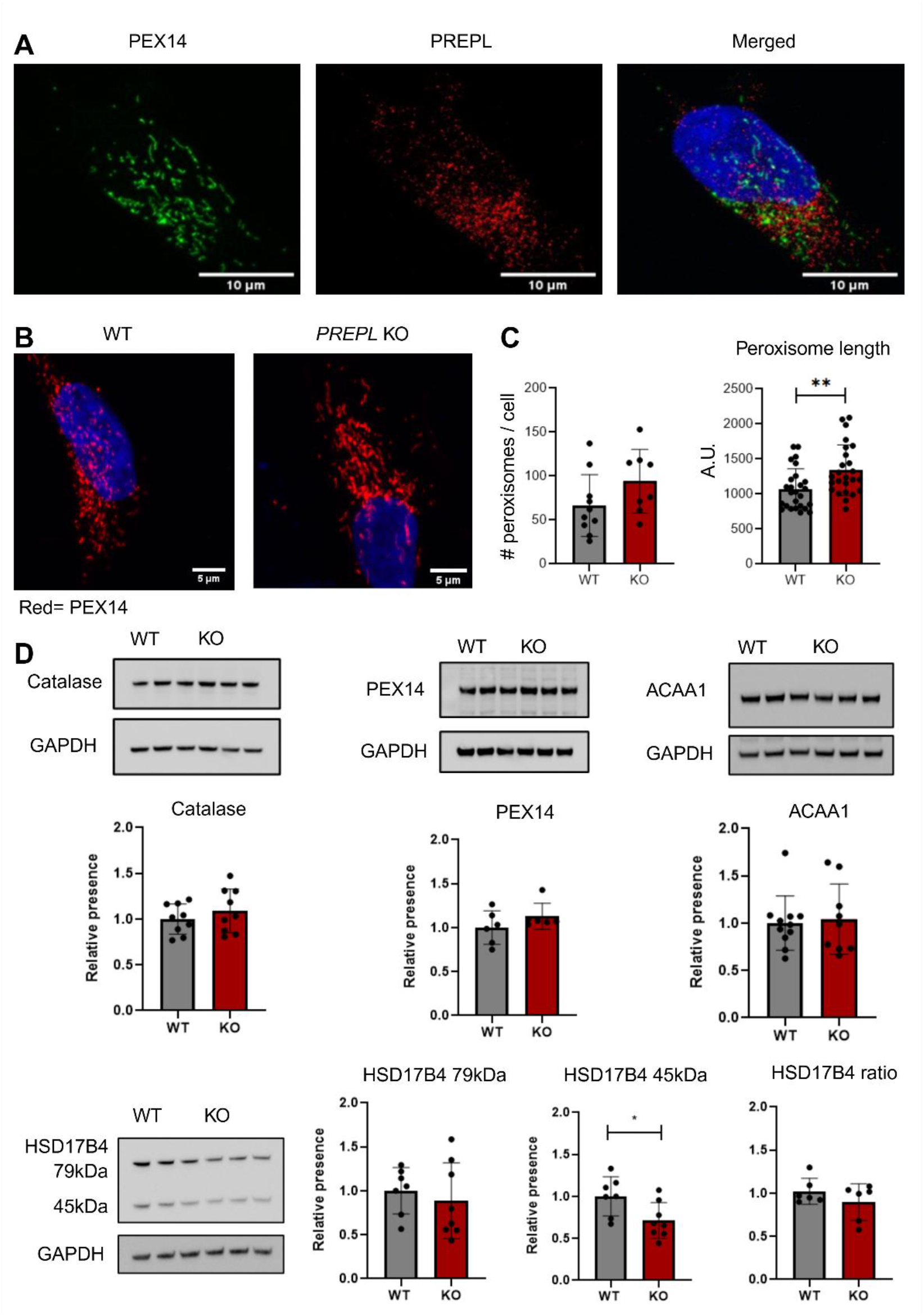
*PREPL* KO alters peroxisome morphology. (A) Co-staining of PREPL (red) with the peroxisomal marker PEX14 (green). The nuclei were counterstained with DAPI (blue) (B) Peroxisome staining in WT and *PREPL* KO HEK293T cells (red: PEX14; blue: DAPI). (C) Quantification of peroxisome number per cell and peroxisome length in WT and *PREPL* KO HEK293T cells. (D) quantitative western blot of peroxisomal proteins in WT and *PREPL* KO HEK293T cells. Statistical analysis was performed using a T-test ^*^p < 0.05, ^**^p < 0.01.

## Discussion

In this study, we investigated whether PREPL exerts a direct role in lipid metabolism as a lipase. Across multiple systems, including brain tissue from *Prepl* KO mice and CRISPR-Cas9-engineered KO cell lines, we consistently observed only modest and variable increases in lysophospholipids. These shifts were not statistically significant and lacked consistency across lipid species, indicating that PREPL does not function as a canonical lysophospholipase. It might however suggest a slight increase in membrane remodeling or turnover in the KO condition. Using a different lipidomics approach on HEK293T cells, a more robust phenotype emerged. *PREPL* KO cells have a significant accumulation of triacylglycerols (TAGs) and an increase in lipid droplet number. This selective lipid storage phenotype is unlikely to reflect a primary lipid-metabolizing role of PREPL. Instead, it points toward perturbations in mitochondrial metabolism. Triacylglycerol accumulation is a common adaptive response to impaired mitochondrial fatty acid oxidation: by sequestering excess fatty acids into neutral lipids, cells protect against lipotoxicity and oxidative stress (18). Direct knockouts of OXPHOS subunits in mitochondria have been shown to lead to an increase in lipid droplets (19). Consistent with this view, mitochondria are known to exert tight control over the balance between lipid oxidation and storage. Thus, the most likely explanation for our findings is that the observed lipidomic alterations represent secondary adaptations to mitochondrial defects.

Indeed, previous studies have shown that PREPL localizes to mitochondria and that its absence results in profound structural and functional abnormalities (4,5,7). *PREPL* KO cells and tissues exhibit reduced cristae density, impaired respiratory chain activity, and reduced respiratory capacity. These defects compromise oxidative phosphorylation capacity and ATP production, while also shifting cellular metabolism toward glycolysis. Moreover, PREPL deficiency possibly leads to changes in ROS through electron leakage and superoxide production caused by a dysfunctional respiratory chain. From a lipid perspective, such mitochondrial dysfunction has two major consequences: (i) impaired β-oxidation, which reduces the catabolism of fatty acids, and (ii) altered NAD^+^/NADH homeostasis, which limits the redox balance required for lipid utilization. Together, these impairments drive the accumulation of fatty acids, which are then diverted into TAG synthesis and stored in lipid droplets as a protective mechanism (18). Thus, the increase in lipid droplet number observed in *PREPL* KO cells likely reflects a direct downstream consequence of defective mitochondria.

Our analysis of peroxisomes further supports this interpretation. PREPL does not localize to peroxisomes, and the levels of canonical peroxisomal proteins remain unchanged in KO cells. Nevertheless, PREPL deficiency leads to elongated peroxisomes. Analogous to findings in the methylotrophic yeast *Pichia pastoris*, it is tempting to speculate that this morphological change may be a consequence of mitochondrial dysfunction (20). Peroxisomes and mitochondria are metabolically interdependent organelles: peroxisomes shorten very-long-chain fatty acids, which are subsequently oxidized in mitochondria, however little is known about how defects in one organelle may affect the other (21).

Taken together, this study provides the first molecular insight into PREPL’s role in lipid metabolism. Rather than acting directly on (lyso)phospholipids or peroxisomal substrates, PREPL deficiency appears to disrupt mitochondrial integrity and function, with minor downstream consequences on cellular lipid balance. Elevated TAG levels and altered peroxisomal morphology likely represent adaptive strategies to maintain lipid and energy homeostasis under conditions of mitochondrial stress. Importantly, these results align with emerging evidence that PREPL is a mitochondrial protein required for cristae architecture, respiratory efficiency, and energy balance.

Future work should prioritize direct assessments of mitochondrial metabolism in *PREPL* KO systems, including measurements of fatty acid oxidation rates and redox balance. Moreover, tracing experiments with stable isotope-labeled fatty acids could clarify whether increased TAGs in PREPL KO cells arise from reduced β-oxidation or altered lipid trafficking. Finally, given the cooperative interplay of mitochondria, peroxisomes, and lipid droplets, it will be important to dissect how PREPL deficiency perturbs organelle communication and whether this contributes to the neuromuscular and metabolic phenotypes associated with CMS22.

